# Ultra-low input single tube linked-read library method enables short-read NGS systems to generate highly accurate and economical long-range sequencing information for *de novo* genome assembly and haplotype phasing

**DOI:** 10.1101/852947

**Authors:** Zhoutao Chen, Long Pham, Tsai-Chin Wu, Guoya Mo, Yu Xia, Peter Chang, Devin Porter, Tan Phan, Huu Che, Hao Tran, Vikas Bansal, Justin Shaffer, Pedro Belda-Ferre, Greg Humphrey, Rob Knight, Pavel Pevzner, Son Pham, Yong Wang, Ming Lei

## Abstract

Long-range sequencing information is required for haplotype phasing, *de novo* assembly and structural variation detection. Current long-read sequencing technologies can provide valuable long-range information but at a high cost with low accuracy and high DNA input requirement. We have developed a single-tube Transposase Enzyme Linked Long-read Sequencing (TELL-Seq^™^) technology, which enables a low-cost, high-accuracy and high-throughput short-read next generation sequencer to routinely generate over 100 Kb long-range sequencing information with as little as 0.1 ng input material. In a PCR tube, millions of clonally barcoded beads are used to uniquely barcode long DNA molecules in an open bulk reaction without dilution and compartmentation. The barcode linked reads are used to successfully assemble genomes ranging from microbes to human. These linked-reads also generate mega-base-long phased blocks and provide a cost-effective tool for detecting structural variants in a genome, which are important to identify compound heterozygosity in recessive Mendelian diseases and discover genetic drivers and diagnostic biomarkers in cancers.

---

Many second-generation sequencing technologies, which sequence thousands to millions of templates at once, have been developed since 2005^1-4^. They can generate tera-bases of highly accurate sequencing output in a run and are used widely in laboratories today. However, their short read length (150 to 600 bp) limits their ability to resolve haplotypes, assemble complex genomes and detect structural variants. Third-generation sequencing platforms that use single-molecule sequencing and promise long-read sequencing capability have also been on the market for nearly a decade, such as SMRT-sequencing^5^ and nanopore sequencing^6^. However, they still yield lower sequencing accuracy at higher sequencing cost than second-generation platforms, and require micrograms of DNA for library construction, which is very challenging to obtain for many real-world samples or applications.

One (sequencing) system for all (applications) is the desire of the customer but requires a significant breakthrough of an enabling technology on either second- or third-generation sequencing platforms. In the past decade, numerous methods have been developed to capture long-range information with short sequencing reads, including mate-pair^7,8^, clonal-barcoding methods (e.g. synthetic long reads^9-11^, linked-reads^12-14^) and Hi-C^15^. Of these, clonal-barcoding library technologies^9-14^ showed the most promising results to bring routine long-read capability to second-generation platforms. The general concept underlining these clonal-barcoding technologies is to uniquely label sub-fragments of a long genomic DNA fragment with a common barcode sequence when the long fragment breaks into small sub-fragments, which later become the inserts of a sequencing library. These methods can be classified into two categories, synthetic long read (SLR) and linked-reads. SLR methods, e.g. LFR^9^, LRseq^10^ and TSLR^11^, in principle have the ability to assemble those small sub-fragments back into their original long fragment with the barcode information but only work for limited fragment lengths (up to 10 Kb) and face technical challenges and expensive operational costs to scale up for large numbers of samples. Linked-reads, sometimes called cloud reads or sparse SLR, have limited potential to re-assemble barcode reads from sub-fragments back into their original long fragment completely, but have been demonstrated to successfully phase the haplotypes, assemble *de novo* genomes and detect structural variants with significantly easier sample preparation^12-14^. Adoption of current linked-read methods is constrained by their dependency on costly instruments for droplet partition^12^, complicated workflows^12-14^, limitations on genome size^12-14^ or incompatibility with widely used sequencing platforms^14^.

To enable a routine linked-read sequencing protocol on a second-generation sequencer in all laboratories for all users, we developed an ultra-low input single-tube linked-read library method, Transposase Enzyme Linked Long-read Sequencing (TELL-Seq^™^). The TELL-Seq method enables barcoding as little as 0.1 ng of genomic DNA in a single PCR tube with 3-hour library construction, without any dedicated specialized instrument.

Both Tn5 and MuA transpososomes have been previously used to simultaneously fragment DNA and introduce adaptors *in vitro,* creating sequencing libraries for next-generation DNA sequencing^16,17^. These protocols remove any long-range information as a result of breaking the long DNA molecules into untraceable small fragments. However, when transpososomes attack DNA targets, they form strand transfer complexes (STCs) which are very stable under natural conditions^18-22^. Only under harsh conditions, such as heat, protease or SDS treatment *in vitro,* will these STCs be disassembled, breaking DNA targets into tagged fragments. This unique feature of the transposition reaction has been used to clonally barcode tagging fragments using Tn5 system^13,14^. One method is to anchor the Tn5 transposon and transposase on a barcoded solid bead surface, then react with genomic DNA^13^; another method is to have the Tn5 transposon and transposase react with genomic DNA first, then ligate them to barcoded beads^14^. We tested both methods using a MuA transposition system. However, neither method was efficient (Supplementary F
ig. 1). We speculated that when both transposon and transposase was immobilized on the bead surface as the first method, their fixed location and spatial arrangement would restrain the efficiency of capturing the free-floating DNA targets by strand transfer reaction only. The limitation for the latter method was that a DNA target full of STCs after strand transfer reaction would create significant steric hindrance from the tertiary structure of protein and DNA complex, and reduce their chance of being captured onto the bead surface. The TELL-Seq method overcame the limitations of both methods with simultaneous strand transfer and transpososome capture reactions in the same solution (Fig. 1a). Barcoding reaction efficiency was significantly improved when both strand transfer and hybridization reactions were dynamically used to capture the DNA targets (Supplementary Fig. 1b). In order to keep the barcoding reaction clonal, we also used non-barcoded beads as a spacer between clonal barcoded beads and increased the solution viscosity to slow DNA diffusion and keep the beads suspended. The TELL-Seq molecular barcode located at the index 1 position in the TELL-Seq library (Fig. 1b) is comprised of 18 degenerate nucleotides with a maximum homopolymer length of six bases and over 2.4 billion unique barcodes. This vast barcoding capability enables the TELL-Seq library method to assign one unique barcode to a single DNA target for maximum barcoding resolution.

**Figure 1.**
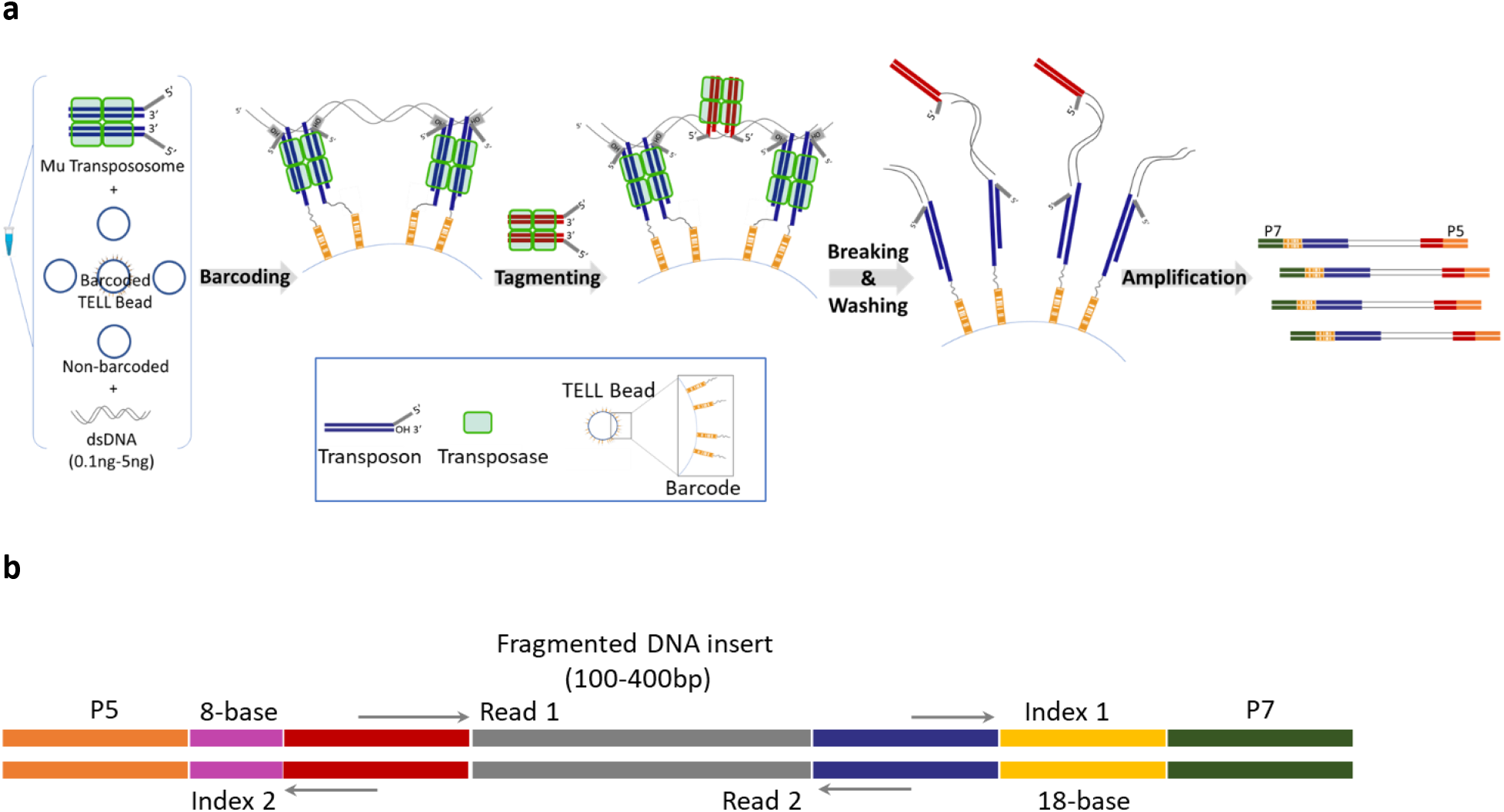
Overview of TELL-Seq library workflow and structure. (**a**) Diagram of TELL-Seq library preparation procedure. In a 0.2 ml PCR tube, 0.1 ng to 5ng genomic DNA based on the genome size were mixed with 3 to 10 million of barcoded TELL Beads and transpososomes for the clonal barcoding reaction. Genomic DNA fragments were captured on the barcoded TELL Beads via connecting strand transfer complexes (STCs) to barcode oligos on the bead surface. A tagging between STCs by a second transpososome introduced a second priming site for library amplification. After breaking the STCs and washing the magnetic TELL Beads, sequencing library molecules were amplified off beads with P5 and P7 adaptor sequences incorporated at the same time. Total library procedure takes approximately three hours. (**b**) TELL-Seq library structure for Illumina sequencing systems. Index 1 location comprises 18-base TELL-Seq molecular barcode; Index 2 location comprises 8-base sample barcode for sample indexing.

We first evaluated the TELL-Seq library method for *de novo* sequencing of microbes. One challenge for microbial sequencing is that the amount of genomic DNA material is often low for fastidious organisms and environmental samples. We developed an ultra-low input TELL-Seq protocol, which used 0.1 ng to 0.5 ng of genomic DNA from 1 Mb to 50 Mb size microbial genomes for library construction. Several bacterial samples were tested with this protocol (Table 1). For *Escherichia coli* DH10B, either 0.1 ng or 0.5 ng genomic DNA were used with three million barcoded TELL beads for the DNA barcoding reaction and approximately one million or 0.2 million reacted TELL beads were used for amplification to generate a paired-end library for 2×146 paired-end sequencing on an Illumina sequencing system, respectively. In order to effectively use TELL-Seq linked-read data for microbial *de novo* assembly, we developed a new *de novo* genome assembler, TuringAssembler, a de Bruijn graph-based assembler that uses linked-read information to perform local assembly and scaffolding in order to produce high-quality assemblies. TuringAssembler works very well for small genome assembly. With the optimal k-mer condition, we achieved excellent assembly results for a set of ultra-low input *E. coli* DH10B samples (Table 1 and Supplementary Table 1). Assembly results from 0.1 ng input were even better than those from 0.5 ng input based on the largest alignment length and the number of misassemblies. This could be due to lower genomic DNA molecule to barcoded bead ratio in the 0.1 ng input condition, which in turn decreased the number of different genomic DNA inputs sharing the same barcode and reduced the ambiguity of linked-read clonality during the assembly process. We sequenced additional bacteria including those with different GC content in the genome, such as *E. coli* K12 MG1655, *Campylobacter jejuni,* and *Rhodobacter sphaeroides* (Table 1 and Supplementary Table 1). The genome of *E. coli* K12 MG1655, lacking the 113,260 bp tandem duplication and many insertion sequences found in *E. coli* DH10B^23^, was assembled better than that of DH10B with 99.9% of the genome assembled with zero misassemblies. *R. sphaeroides* with 68.8% GC content was a more challenging genome to sequence and assemble. Its assembly results showed many small contigs with length less than 5000 bp. The genomic DNA of *R. sphaeroides* was purchased and extracted with a standard method having average length of just 20 Kb; while genomic DNA from both strains of *E. coli* was prepared with a high molecular weight specific protocol and averaged over 40 Kb in length. The shorter genomic DNA length of *R. sphaeroides* may have also contributed to lower assembly performance compared with the *E. coli* samples.

**Table 1.**
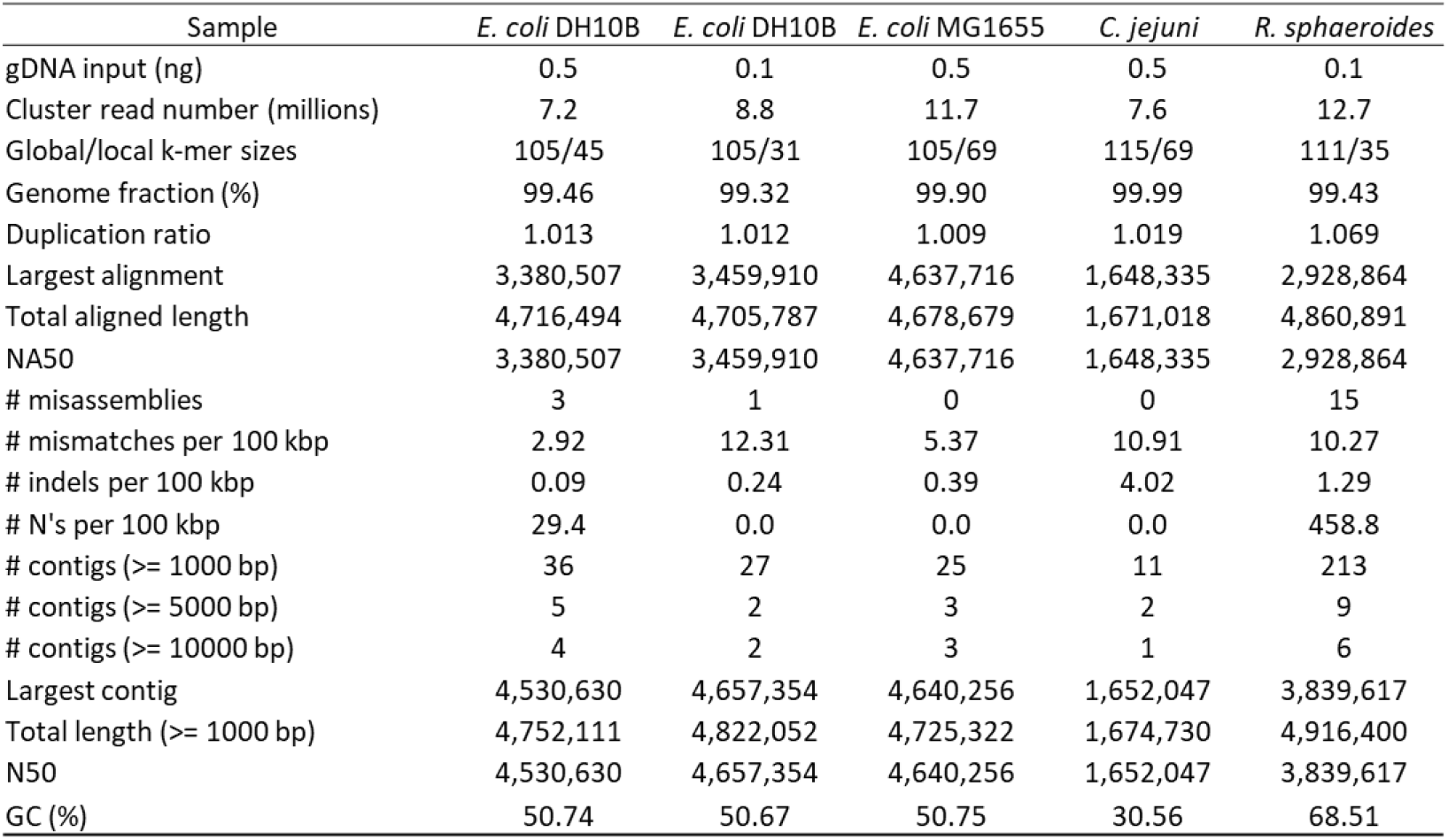
Summary of *de novo* assembly results using TuringAssembler on microbial samples.

Furthermore, we sequenced and assembled the genome of a human gut microbial isolate, *Coprobacillus cateniformis* DSM-15921 (Firmicutes), using 0.1 ng genomic DNA. Most of the contigs generated from the assembly aligned well to *C. cateniformis,* as expected. There were seven assemblies previously deposited for this organism in the NCBI database. The best assembly among them was AKCB01, which was a hybrid assembly from Illumina short reads and PacBio long reads. In addition, the whole genome shotgun (WGS) assembly CABKQT01 was identical to AKCB01 when comparing their assembly results and was excluded from further evaluation. Our assembly results for DSM-15921 (Supplementary Table 2 and 3) showed the longest N50 contig length (3,607 Kb) among all previously deposited assemblies for this organism and had the highest ratio of largest contig length to total length, 98.4%. To quantitively assess the assembly quality, we ran a BUSCO analysis using Firmicutes as the bacterial lineage^24^ and identified 221 complete and single-copy BUSCO groups out of 232 expected BUSCO groups with 11 BUSCO groups missing in this lineage (Supplementary Table 4). The 95% complete rate of the DSM-15921 assembly was among the highest of all assemblies. We further confirmed that the 11 BUSCO groups missing in our assembly were absent in all six deposited assemblies (Supplementary table 4).

TELL-Seq is the first linked-read technology demonstrated for efficient microbial genome sequencing and the first long-range sequencing technology enabled under nanogram input whole genome sequencing. Other linked-read methods have been effectively applied to whole genome haplotype phasing and structural variation detection for human samples^12-14^. With over two billion unique barcodes, the TELL-Seq technology should be easily used for such applications as well. As a demonstration for such human applications, we applied TELL-Seq method to two well-characterized Genome in a Bottle (GIAB) consortium samples NA12878 and NA24385. Five nanograms of each input DNA were processed to construct TELL-Seq libraries with approximately eight million barcoded TELL Beads, then pooled and sequenced using a S1 flowcell on a NovaSeq 6000 with 2×146 paired-end reads workflow. Data were analyzed as described in the Methods section and results were reported in Table 2. The HapCUT2 phasing tool was used for haplotype phasing^25^. The NovaSeq run generated 1,024 million and 959 million cluster reads for NA12878 and NA24385 samples (total 1,983 million) from a single S1 flowcell run, respectively (Table 2). Greater than 96% of reads from both samples were mapped to the GRCh38 reference. There were approximately 7.7 million and 7.4 million effective barcodes identified for the two samples, respectively. Over 90% linked-read molecules had lengths greater than 20 Kb, whereas 20-30% linked-read molecules had lengths greater than 100 Kb. Over 99.8% of the heterozygous single nucleotide variants (SNVs) were phased in each sample with N50 phasing block size greater than 16 Mb and low switch error rates (Table 2). The manufacturer’s sequencing throughput specification for a S1 flowcell is 1,300 million to 1,600 million cluster reads and there are a 200-cycle and a 300-cycle sequencing chemistry available for this flowcell. We subsampled our sequencing reads for each sample down to under 700 million each and 2×95 read length to simulate a lower loading density on the S1 flowcell run with 2×96 paired-end workflow using a 200-cycle kit. With the subsampled read number and read length, we were still able to phase 99.7 % heterozygous SNVs in each sample with N50 phasing block size greater than 7.7 Mb while keeping low switch error rates (Table 2). Phasing results from HapCUT2 analysis outperformed those from Long Ranger^™^, which is also a widely used phasing tool and designed for 10x linked-read specifically. For the same NA12878 TELL-Seq dataset, Long Ranger analysis showed approximately 98.9% heterozygous SNVs were phased with N50 phasing block size at 4.2 Mb only.

**Table 2.** Summary of TELL-Seq phasing results on NA12878 and NA24385 samples.

| Sample | NA12878 | NA24385 | NA12878-Trimmed<br>To 95 bases;<br>subsample <700M | NA24385-Trimmed<br>To 95 bases;<br>subsample <700M |
| --- | --- | --- | --- | --- |
| Barcodes | 7,711,261 | 7,389,133 | 7,280,000 | 7,033,277 |
| Sequencing condition | 2x146 PE | 2x146 PE | 2x96 PE* | 2x96 PE* |
| Cluster read number (millions) | 1024 | 959 | 696 | 695 |
| Mapped % | 97.3% | 97.7% | 96.3% | 96.8% |
| Duplicates % | 43.1% | 35.0% | 35.4% | 28.9% |
| Mean depth of coverage (X) | 77.2 | 74.1 | 38.8 | 39.2 |
| Mean depth of coverage (duplicates removed) | 43.9 | 48.2 | 25.1 | 27.9 |
| Mean DNA/TELL bead (Kb) | 199.9 | 245 | 187.6 | 226.3 |
| DNA in molecules >20kb: | 93.2% | 90.0% | 93.2% | 90.1% |
| DNA in molecules >100kb: | 33.9% | 20.6% | 32.7% | 19.9% |
| Weighted mean molecule length (kb) | 46 | 36.7 | 45.7 | 36.7 |
| N50 Reads per molecule | 44 | 28 | 36 | 24 |
| hetSNVs phased (%) | 99.9 | 99.8 | 99.7 | 99.7 |
| Phasing block N50 (Mb) | 16.1 | 17.5 | 7.7 | 9.4 |
| Longest phasing block (Mb) | 67.5 | 59.2 | 39.9 | 35 |
| Short switch error rate (%) | 0.041 | 0.075 | 0.065 | 0.133 |
| Long switch error rate (%) | 0.036 | 0.081 | 0.046 | 0.118 |

\* Simulation of 2x96 paired-end run with 200-cycle sequencing kit at lower loading density on a S1 flowcell by trimming back the sequencing read length and subsampling total sequencing reads.

We also checked the variant calling results without incorporating any barcode information on the NA12878 sequencing data and obtained 99.1% recall rate and 98.9% precision rate on SNVs and 89.8% recall rate and 89.3% precision rate for INDEL variants against the GIAB high-confidence benchmark variant calls with filtering conditions described in the Methods section (Supplementary Table 5).

Previous studies have reported ten structural variants (i.e., deletions) in NA12878 revealed by other linked-read methods^12-14^ (Fig. 2a). However, there are discrepancies regarding two of these SV calls between the 10x linked-read method^12^ and stLFR method^14^. With respect to the deletion at chr.3: 162512134-162626335, Zhang et al (10x) identified it as a 114 Kb heterozygous deletion^12^, whereas Wang et al (stLFR) claimed it as a 19 Kb homozygous deletion^14^. We used Long Ranger for phasing and structural variation detection and Loupe^™^ for visualization of TELL-Seq data. Our data (Fig. 2b and 2c) clearly resolved this SV as a small 19 Kb homozygous deletion within a large 114 Kb heterozygous deletion, which could explain why other linked-read methods called it differently. With respect to the deletion at chr.5: 104431113-104503673, Zhang et al identified it as a heterozygous deletion^12^, while Wang et al claimed it as a homozygous deletion^14^. Our data provided support to classify it as a heterozygous deletion (Supplementary Fig. 6). For this case, a heterozygous deletion call was more reliable than a homozygous deletion call which could be due to lack of sequencing coverage rather than an actual deletion. We also confirmed the other eight SVs in the TELL-Seq data as heterozygous deletions (Supplementary Fig. 3 to Fig. 10) as reported previously by others.

**Figure 2.**
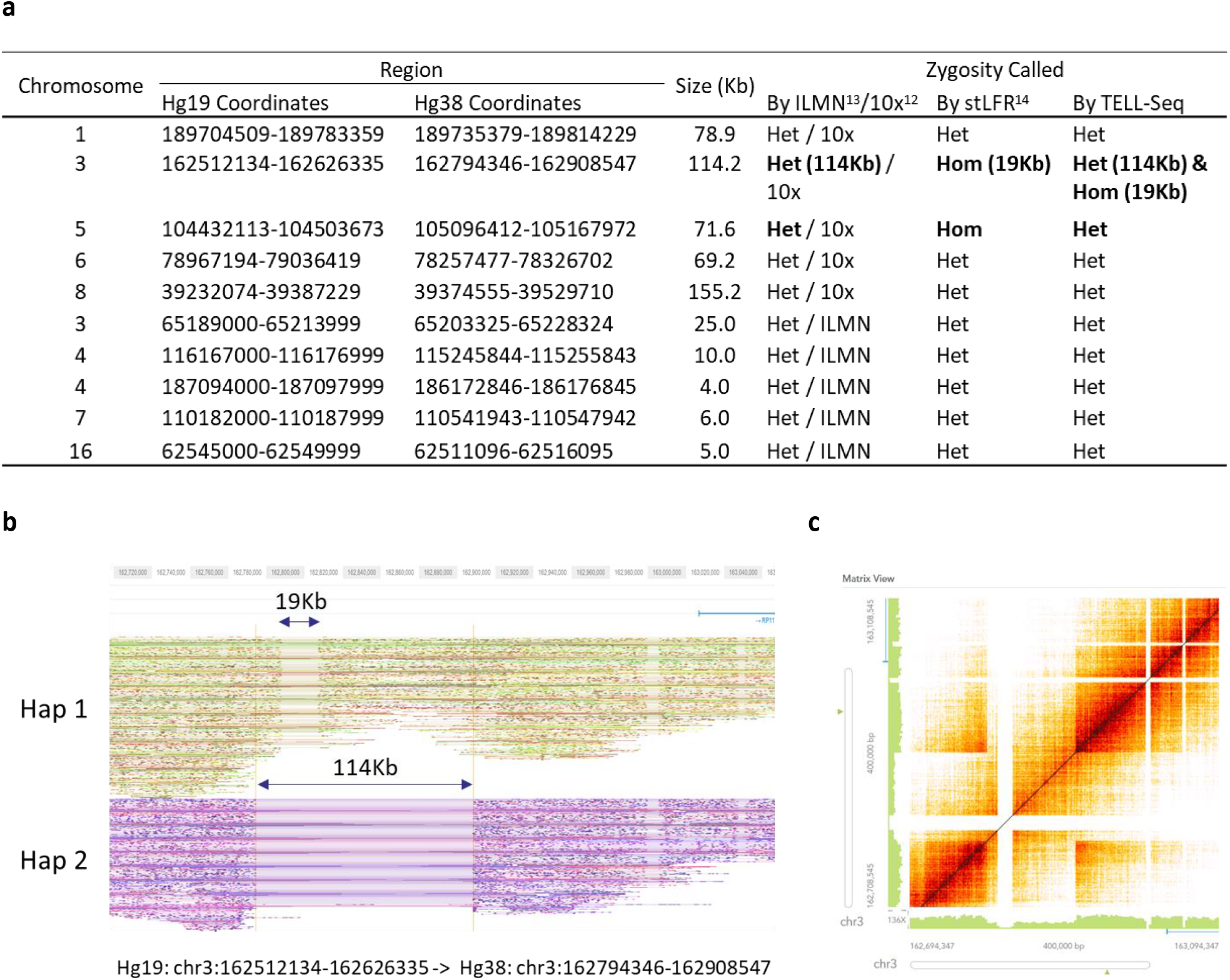
Detection of structural variations in NA12878. (**a**) Comparison of ten large deletion calls reported by different linked-read methods from 10x^12^, Illumina’s CPTv2-seq^13^, and stLFR^14^. (**b**) Phased read graph from TELL-Seq data showed a 19 Kb homozygous deletion (Hom) within a 114 Kb heterozygous deletion (Het) on chromosome 3: 162512134-162626335. For the same location, 10x data^12^ only identified a 114 Kb heterozygous deletion, while stLFR data^14^ only identified a 19 Kb homozygous deletion. Hg38 coordinates were used for visualization data. (**c**) Heatmap of the same region from TELL-Seq data clearly showed the presence of both the small homozygous deletion and the large heterozygous deletion.

In addition, we generated a *de novo* assembly with the TELL-Seq data for NA12878 using SuperNova 2.1.1. Although the TELL-Seq library insert size averaged 200 bp and was much shorter than the optimal condition required for SuperNova (350-400 bp), we used default parameters for the SuperNova analysis except with reads longer than 125 bp for both R1 and R2. N50 and NA50 scaffold length were 31.5 Mb and 4.3 Mb, respectively. Largest contig and largest alignment lengths were 109.2 Mb and 23.6 Mb, respectively (Supplementary Table 6). In comparison to assembly results using other linked-read methods^12,14^ and nanopore long-read sequencing^26^, the TELL-Seq derived assembly showed longer aligned contig length and at least 28% and 71% fewer misassemblies than other linked-read methods and nanopore method, respectively (Supplementary Table 7).

Here we demonstrated TELL-Seq as a streamlined linked-read technology for whole genome sequencing of a variety of genome sizes and could reliably detect SVs that were misclassified by other linked-read methods. Recently, 10x linked-reads were used for diplotyping of CRISPR-Cas9 captured targeted gene loci ranging from the 200 Kb BRAC1 gene to the 4 Mb MHC locus^27^. These target sizes fit well with our TELL-Seq technology as our ultra-low input protocol for small bacterial genome sequencing has demonstrated. In addition, with capacity of over two billion unique barcodes, the TELL-Seq technology can assign one unique barcode to each single large DNA molecule, which will be critical for application of phasing small targeted gene loci.

For human SV detection and *de novo* assembly we used analysis tools developed for the 10x linked-read method specifically. This worked for TELL-Seq data in general. However, due to the short library insert length and different barcoding chemistry of TELL-Seq, these tools are not optimized for TELL-Seq data. Further fine tuning and data training of these tools should improve the sensitivity of SV calls and reduce the proportion of short contigs in a *de novo* assembly.

We present an easy-to-use and easy-to-automate single-tube linked-read library method that can cost-effectively generate long-range information from short-read NGS systems. The lengths of many linked molecules generated from sub-nanogram to nanogram input material are over 100 Kb. Such long-range information generated from such low input is very difficult to achieve by current commercially available long-read sequencing technologies. With the TELL-Seq library technology, a routine linked-read library for whole genome and contiguous targets will become a reality for accurate haplotype-resolved sequencing and *de novo* sequencing.

## Methods

### Genomic DNA

Genomic DNA of *C. jejuni* and *R. sphaeroides* were purchased from ATCC and used directly without any size selection. Average genomic DNA size of *C. jejuni* and *R. sphaeroides* were 28Kb and 20Kb, respectively.

Genomic DNA of *E. coli* DH10B and K12 MG1655 were extracted using a modified salting-out protocol^28^ described below. NA12878 and NA24385 DNA were extracted from harvested immortalized human lymphocyte cells GM12878 and GM20847 (Coriell Institute, Camden, NJ) using the same salting-out protocol, respectively. Briefly, 5×10^6^ human/bacterial cells were resuspended in 3 ml of 10 mM Tris, 400 mM NaCl, 2 mM EDTA, pH 8.0 and lysed by the addition of 0.2 ml 10 % SDS and 0.5 ml Proteinase K solution [1mg/ml Proteinase K (Ambion, Austin, TX), 1 % SDS, 2 mM EDTA, pH 8.0]. After an overnight incubation at 37 °C (12-18 hours), the cell lysate was mixed with 1.2 ml of 5 M NaCl and centrifuged at 1100 g for 15 min at 4 °C. The supernatant was transferred and mixed with 8 ml of 100% ethanol and centrifuged at 8000 g for 15 min at 4°C to precipitate the DNA. The pellet was air-dried and resuspended in 50 μl TE. Following an incubation with 20 μg RNase A (Thermo Fisher Scientific, Waltham, MA) at room temperature for 30 min, genomic DNA was stored at 4°C in a DNA low-bind tube.

*C. cateniformis* DSM-15921 was obtained from DSMZ culture collection and grown inside of a vinyl anaerobic chamber (Coy Laboratory Products, Grass Lake, MI) with an atmosphere of 3% hydrogen, 10% CO2 and nitrogen as balance. Liquid pure cultures were grown in anoxic BHI broth supplemented with hemin, vitamin K and L-cysteine. Genomic DNA of strain DSM-15921 was extracted from pure culture using the MagMAX-96 DNA Multi-Sample Kit (Applied Biosystems, Foster City, CA). We followed the manufacturer’s instructions for isolating genomic DNA from cultured cells, but extracted in 1.5-mL microcentrifuge tubes in place of a 96-well plate, and used a Hula Mixer (Life Technologies, Carlsbad, CA) in place of a titer plate shaker. For the elution step, we followed instructions for the non-heated shaking option. After extraction, no vortexing was applied and only wide-boar tips were used to facilitate recovery of high molecular weight DNA fragments. We then performed a 0.4X AMPure size-selection and bead clean-up (Beckman Coulter, Brea, CA), followed by further size selection with a Short-Read Elimination (SRE) kit (Circulomics, Baltimore, MD), which removes all fragments < 10 Kb. Size-selected genomic DNA was stored at 4°C in a DNA low-bind tube.

### TELL-Seq library construction and sequencing

TELL-Seq libraries were constructed using a TELL-Seq WGS Library Prep Kit (Universal Sequencing Technology, Carlsbad, CA). Briefly, 0.1 ng or 0.5 ng genomic DNA from microbial samples or 5 ng genomic DNA from human samples were used with approximately 3 million or 8 million TELL beads for barcoding reaction in a 0.2 ml PCR tube according to the manufacturer’s protocol, respectively. TELL beads are 3 μm magnetic beads, each having at least one unique barcode sequence conjugated on the surface. There are approximately 50,000 copies of total barcode templates on each TELL bead. Among them, approximately 1% barcode templates share at least one common barcode sequence with other TELL beads. After the barcoding reaction, TELL beads with captured barcoded DNA were amplified for 13-14 cycles for microbial samples and 8 cycles for human samples to produce final sequencing libraries. The high-throughput nature of the reaction allows for construction of multiple libraries in a 96-well format by one person in 4 hours. The TELL-Seq libraries were quantified by Qubit dsDNA BR Assay kit (Thermo Fisher Scientific, Waltham, MA) and pooled for sequencing on a MiSeq/NextSeq for microbial samples or a NovaSeq for human samples with 2×146 paired-end reads, 18-cycle Index 1 reads and 8-cycle Index 2 reads based on the manufacturer’s protocols.

### Primary sequencing data processing

The sequenced raw data were first processed by the TELL-Read analysis pipeline software for barcode correction and filtering before proceeding to downstream analysis, such as, phasing, variant calling, SV detection, and *de novo* genome assembly.

After sequencing, raw read BCL files were converted to FASTQ files using bcl2fastq and adapter sequences were removed. The FASTQ files were then demultiplexed into index I1 reads, R1 reads and R2 reads for each sample based on the Index 2 reads. I1 reads are the TELL-Seq barcode sequences. For each sequencing library construction, a set of unique barcode sequences were randomly chosen from a 2.4 billion barcode pool. After sequencing, all unique barcodes were identified along with the count for the number of reads they were associated with. The unique barcodes associated with only one read were error corrected if they were 1-base mismatched with one of the barcodes associated with multiple reads. Barcodes with errors after this step were filtered out. The erroneous barcodes along with their associated reads were removed and excluded from the rest of analyses.

### *De novo* assembly of microbial samples

TuringAssembler was developed and used for microbial genome assembly (https://bioturing.com/resources/turingassembler/TuringAssembler_x64_Linux). It combines paired-end read and linked-read barcode information to perform local assembly in order to resolve local complex regions caused by tandem duplications; linked-read information is also applied for scaffolding the contigs resulting from early assembly steps.

The process that makes TuringAssembler unique is local assembly. For two contigs that are predicted to be consecutive on the genome based on both paired-end reads and barcode information, finding the path between them on the assembly graph is not straight forward because they usually are stitched into a complex region, together with other pairs of contigs. This region usually comprises similar k-mer compositions from many copies of a repetitive sequence in the genome. Rather than ignoring it and filling ambiguous characters between the two consecutive contigs, TuringAssembler attempts to *de novo* assemble the repeat region between them locally using linked-read information.

First, reads that originated from the start and the end of those contigs along with reads within the gap region between the contigs are isolated using barcode information. We denote the barcode tagged on a read *r* as *b*(*r*). *R*(*x*) denotes the set of reads that can be aligned on the region *x*. The set of barcodes that “span” a region *x* is denoted as *B(x) = {b(r) ∈ r ∈ R(x)}·* A long contig *C* (larger than 4 Kb) has two bounded regions *C^h^*, at the beginning (head) of *C*, and *C^t^*, at the end (tail) of *C*. Each region *C^h/t^* has the length 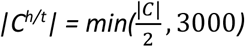. For example, a pair of long contigs *C_1_* and *C_2_* has the representation on the genome as (*C_1_^t^, C_1_^h^, C_2_^t^, C_2_^h^*. The set of barcodes that span *C_1_^h^* or *C_2_^t^ is denoted as* B_union_ = B(*C_1_^h^*) ∪ B(*C_2_^t^*). The set of barcodes that span both *C_1_^h^* and *C_2_^t^ is denoted as* B_share_ = B(*C_1_^h^*) ⋂ B(*C_2_^t^*). The set of reads that is used to construct the local de Bruijn graph for the region (*C_1_^h^, C_2_^t^*) is denoted as *R_union_ = {r/b(r) ∈* B_union_*}*. Since this approach still unable to capture short molecules that are located completely in the gap region, in order to keep the graph connected, we use smaller k-mer size to construct the “local” de Bruijn graph. We next identify a pair of edges *E_1_, E_2_* in the assembly graph *G_local_* that represent the sequences of *C_1_^h^, C_2_^t^*. The nucleotide sequence that “bridges” *C_1_^h^* and *C_2_^t^* can be represented by a path that starts at *E_1_* and ends at *E_2_* in the assembly graph Glocal. There might be multiple paths like that in Glocal. But the true path must represent the true multiplicity of its edges in the local region (*C_1_^h^, C_2t_* and maximize the mapping capability of the local set of reads *R_share_ = {r | b(r) ∈* Bshare *}*. We first use *R_share_* to estimate the coverage of all edges in G_local_ then use the average coverages of *E_1_* and *E_2_* to compute the unit coverage. The multiplicity of each edge in Glocal is the ratio of its coverage to the unit coverage. We use the total number of concordance aligned read-pairs as a score to sort all paths that start at *E_1_* and end at *E_2_*, then choose the nucleotide sequence of the best path as the sequence in the local region *(C_1_^h^, C_2_^t^)*. In case the local region is very complex as we cannot enumerate all paths between *E_1_* and *E_2_*, we fill a series of ‘N’ characters in the region *(C_1_^h^, C_2_^t^)* where the length of the series equals to the shortest path that connects *C_1_^h^* and *C_2_^t^*.

TuringAssembler uses two different k-mer sizes during assembly: a global k-mer size to construct the de Bruijn graph from all input data, and a local k-mer size to construct the de Bruijn graphs in the local regions. The global k-mer size can be inferred from the estimated genome coverage and the read length. It is an odd number close to the estimated genome coverage, but not larger than the average read length. The local k-mer size should be smaller than the global k-mer size and may vary from between datasets. Usually multiple k-mer sizes should be tested in order to identify an optimal k-mer combination for a dataset.

For assembly completeness evaluation, BUSCO^24^ was run using the “-m geno” option to indicate that the input was a genome assembly, using the firmicutes_odb9 lineage as a reference.

### Variant calling and filtering

Paired-end reads with corrected barcode information were mapped to the reference genome using BWA-MEM (0.7.17-r1188), and the mapped BAM file was sorted by chromosome coordinates. After duplicate reads were marked and removed with Picard (v2.18.7), and read group information were added to the BAM file, the germline variants including single nucleotide polymorphisms (SNPs) and small indels between the sample and the reference genome were called using HaplotypeCaller in the GATK tool package (GATK4-4.1.2.0-1). Variants were filtered using the following VCF parameters: QUAL > 15, QUAL < 50, and MQRankSum < 6. SNPs were filtered as those with AD ≥ 2 and AF > 0.15 while short indels had AD ≥ 4 and AF > 0.25. These parameters ensure that the reads were mapped to a unique place in the assembly with high quality, that the reads carrying the alleles were sufficient in terms of frequency, depth, and mapping quality (AD, AF, MQRankSum), and that the actual variants were called with high quality (QUAL). For NA12878 sample, the called and filtered variants were compared with GIAB variant calls (ftp://ftp-trace.ncbi.nlm.nih.gov/giab/ftp/release/NA12878_HG001/latest/GRCh38/) using the Illumina haplotype VCF comparison tool, hap.py (https://github.com/Illumina/hap.py.git).

### Phasing linked-reads using HapCUT2

Linked-reads were phased with HapCUT2 (https://github.com/vibansal/HapCUT2), using heterozygous SNVs that involved two alleles of the same length. The BAM file of diploid variants with duplicates removed was used as input to “extractHAIRS” in the HapCUT2 tool to create the compact fragment file containing only haplotype-relevant information. Linked fragments were generated using the “LinkFragments.py” program in the package. The linked fragments and variants in VCF format were then used as input to “HapCUT2” for phasing.

For assessing the accuracy of the linked-read haplotypes, we used the high-quality phased genotypes for the two individuals, NA12878 and NA24385, from the GIAB project. For NA12878, 99% of the variants are phased using the Platinum Genome pedigree analysis while for the NA24385 genome, 87.0% of the calls are phased using trio analysis. We have previously observed (Bansal, unpublished results) that errors in these haplotypes can inflate the switch error rates when evaluating an independent set of sequencing-based haplotypes. Therefore, we assembled a consensus of the GIAB haplotypes and 10x Genomics linked-read haplotypes (VCF files downloaded from the GIAB ftp site) by discarding the small fraction of variants that are inconsistently phased between the two sets of haplotypes. These consensus haplotypes were used for calculating short and long switch error rates using scripts in the HapCUT2 software package.

### Structural variant detection and *de novo* assembly of human genome

Structural variation detection and visualization were performed using Long Ranger^™^ (v2.2.2) and Loupe^™^ (v2.1.1) tools (10x Genomics, Pleasanton, CA), respectively. First, TELL-Seq data were converted into 10x compatible data format. Briefly, all unique TELL-Seq 18-base barcodes used in each sequencing library were identified and converted to a whitelist of 10x compatible (i.e., 16-base) unique barcodes. The mapping was also done in such a way that any two of the resulting barcodes were 2+ Hamming distance away from each other to avoid error correction step in the Long Ranger process. The converted barcodes followed by 7 ‘N’ characters were then added to the beginning of each corresponding R1 read in FASTQ format. These converted R1 reads, along with R2 reads were used as the input for Long Ranger. In addition, Long Ranger’s barcode whitelist file, e.g., /longranger-2.2.2/longranger-cs/2.2.2/tenkit/lib/python/tenkit/barcodes/4M-with-alts-february-2016.txt was replaced with a newly created TELL-Seq converted barcode whitelist. All subsequent analyses were done following standard Long Ranger procedure. For the human NA12878 sample, to keep total unique barcode count less than 16 million which was the upper limit for the Loupe program, barcodes associated with only one read were removed and Long Ranger v2.2.2 was run with default parameters on the remaining set of 1,014 million cluster reads using the GRCh38-2.1.0 reference and the GATK-3.8-0 variant caller. Results of Long Ranger were visualized with Loupe program v2.1.1. Genome coordinate conversion between GRCh38 (hg38) and GRCh37 (hg19) references was done with Lift Genome Annotations (https://genome.ucsc.edu/cgi-bin/hgLiftOver).

The human genome *de novo* assembly was performed using Supernova (v2.1.1) with default parameters on all reads longer than 125bp on both PE ends.

### Data availability

All sequencing data have been deposited into the NCBI under BioProject PRJNA591637.

## Acknowledgments

We would like to thank Feng Pan for testing data analysis software, Dennis Woods and Alice Tagle for supporting TELL-Seq kit production and Michelle Zhang for assistance with optimizing DNA extraction from DSM-15921. We thank Dr. Wei Wang and the Princeton University Genomics Core Facility for NovaSeq sequencing.

