## Supplemental Tables and Figures for "Ultra-low input single tube linked-read library method enables short-read NGS systems to generate highly accurate and economical long-range sequencing information for *de novo* genome assembly and haplotype phasing"

**Supplementary Table 1. Summary of *de novo* assembly results using TuringAssembler on microbial samples with 5 million or 3 million subsampled sequencing reads from Table 1**

| Sample | *E. coli* DH10B | *E. coli* DH10B | *E. coli* MG1655 | *C. jejuni* | *R. sphaeroides* |
|---|---|---|---|---|---|
| gDNA input (ng) | 0.5 | 0.1 | 0.5 | 0.5 | 0.1 |
| Cluster read number (millions) | 5 | 5 | 5 | 3 | 5 |
| Global/local k-mer sizes | 105/45 | 105/69 | 105/45 | 115/31 | 105/45 |
| Genome fraction (%) | 99.4 | 99.3 | 99.8 | 99.5 | 99.2 |
| Duplication ratio | 1.026 | 1.008 | 1.005 | 1.007 | 1.106 |
| Largest alignment | 2,633,347 | 4,510,002 | 4,618,779 | 1,215,159 | 1,328,898 |
| Total aligned length | 4,772,611 | 4,687,503 | 4,657,114 | 1,644,036 | 5,015,052 |
| NA50 | 2,633,347 | 4,510,002 | 4,618,779 | 1,215,159 | 776,515 |
| # misassemblies | 3 | 2 | 0 | 1 | 20 |
| # mismatches per 100 kbp | 3.69 | 4.3 | 3.37 | 6.12 | 6.98 |
| # indels per 100 kbp | 0.19 | 0.15 | 0.28 | 3.43 | 1.09 |
| # N's per 100 kbp | 84.6 | 52.4 | 9.5 | 0.0 | 510.2 |
| # contigs (>= 1000 bp) | 80 | 23 | 20 | 4 | 355 |
| # contigs (>= 5000 bp) | 5 | 3 | 3 | 2 | 11 |
| # contigs (>= 10000 bp) | 4 | 2 | 3 | 1 | 7 |
| Largest contig | 4,478,097 | 4,626,772 | 4,619,821 | 1,630,556 | 2,837,907 |
| Total length (>= 1000 bp) | 4,802,032 | 4,811,224 | 4,696,118 | 1,639,751 | 5,043,831 |
| N50 | 4,478,097 | 4,626,772 | 4,619,821 | 1,630,556 | 2,837,907 |
| GC (%) | 50.71 | 50.75 | 50.76 | 30.55 | 68.5 |

**Supplementary Table 2. *De novo* assembly results using TuringAssembler on DSM-15921 sample**

| | |
|---|---|
| Global/local k-mer sizes | 105/35 |
| # contigs | 30 |
| # contigs (>= 0 bp) | 30 |
| # contigs (>= 1000 bp) | 20 |
| # contigs (>= 5000 bp) | 3 |
| Largest contig | 3,607,461 |
| Total length | 3,666,615 |
| Total length (>= 0 bp) | 3,666,615 |
| Total length (>= 1000 bp) | 3,659,685 |
| Total length (>= 5000 bp) | 3,622,804 |
| N50 | 3,607,461 |
| GC (%) | 31.38 |
| # N's per 100 kbp | 5.45 |

**Supplementary Table 3. Comparison of *C. cateniformis* assemblies deposited in NCBI database with TELL-Seq results (DSM-15921)**

| Strain/Assembly | Contigs | N50 (Kb) | L50 | Total length (Kb) |
|---|---|---|---|---|
| 29_1/ADKX01 | 80 | 120 | 11 | 3,857 |
| D6/AKCB01 | 9 | 2,786 | 1 | 3,861 |
| JCM/BBDT01 | 101 | 91 | 10 | 3,585 |
| OM02-34/QSVJ01 | 100 | 91 | 12 | 3,755 |
| OM02-5/QSVG01 | 96 | 86 | 12 | 3,750 |
| OF01-6/QSCY01 | 108 | 81 | 15 | 3,803 |
| DSM-15921 | 30 | 3,607 | 1 | 3,666 |

**Supplementary Table 4. BUSCO analysis results using Firmicutes as the bacterial lineage on all the available *C. cateniformis* genome assemblies**

| Assembly | DSM-15921 | ADKX01 | AKCB01 | BBDT01 | QSCY01 | QSVG01 | QSVJ01 |
|---|---|---|---|---|---|---|---|
| Complete BUSCO | 221 | 219 | 221 | 183 | 221 | 221 | 221 |
| Complete and single-copy BUSCOs | 221 | 216 | 221 | 182 | 218 | 218 | 218 |
| Complete and duplicated BUSCOs | 0 | 3 | 0 | 1 | 3 | 3 | 3 |
| Fragmented BUSCOs | 0 | 0 | 0 | 29 | 0 | 0 | 0 |
| Missing BUSCOs | 11 | 13 | 11 | 20 | 11 | 11 | 11 |
| Total BUSCO groups searched | 232 | 232 | 232 | 232 | 232 | 232 | 232 |
| % Complete | 95.3% | 94.4% | 95.3% | 78.9% | 95.3% | 95.3% | 95.3% |

**Supplementary Table 5. Summary of TELL-Seq SNV and Indel variant calling results compared with GIAB reference on NA12878 sample**

| Variant Type | Analysis | Recall Rate | Precision Rate |
|---|---|---|---|
| SNV | No Filter | 99.2% | 97.7% |
| | Filtered | 99.1% | 98.9% |
| INDEL | No Filter | 89.8% | 79.9% |
| | Filtered | 89.8% | 89.3% |

**Supplementary Table 6. QUAST[29] analysis of *de novo* assembly of NA12878 from TELL-Seq data.**

| Assembly | TELL-Seq* |
|---|---|
| # contigs (>= 0 bp) | 34,085 |
| # contigs (>= 1000 bp) | 34,085 |
| # contigs (>= 5000 bp) | 13,584 |
| # contigs (>= 10000 bp) | 7,508 |
| # contigs (>= 25000 bp) | 1,181 |
| # contigs (>= 50000 bp) | 542 |
| Total length (>= 0 bp) | 2,979,445,840 |
| Total length (>= 1000 bp) | 2,979,445,840 |
| Total length (>= 5000 bp) | 2,929,255,010 |
| Total length (>= 10000 bp) | 2,886,528,456 |
| Total length (>= 25000 bp) | 2,794,100,219 |
| Total length (>= 50000 bp) | 2,772,210,571 |
| # contigs | 34,085 |
| Largest contig | 109,183,970 |
| Total length | 2,979,445,840 |
| Reference length | 3,099,922,541 |
| GC (%) | 40.91 |
| Reference GC (%) | 40.87 |
| N50 | 31,462,027 |
| NG50 | 29,617,959 |
| N75 | 12,845,396 |
| NG75 | 10,385,879 |
| L50 | 28 |
| LG50 | 30 |
| L75 | 65 |
| LG75 | 73 |
| # misassemblies | 1,987 |
| # misassembled contigs | 939 |
| Misassembled contigs length | 2,677,809,708 |
| # local misassemblies | 1,549 |
| # scaffold gap ext. mis. | 9,216 |
| # scaffold gap loc. mis. | 20,150 |
| # unaligned mis. contigs | 74 |
| # unaligned contigs | 4158 + 1220 part |
| Unaligned length | 23,049,908 |
| Genome fraction (%) | 93.732 |
| Duplication ratio | 1.075 |
| # N's per 100 kbp | 6419.8 |
| # mismatches per 100 kbp | 114.07 |
| # indels per 100 kbp | 25.61 |
| Largest alignment | 23,573,913 |
| Total aligned length | 2,771,334,486 |
| NA50 | 4,302,918 |
| NGA50 | 4,086,890 |
| NA75 | 1,495,528 |
| NGA75 | 1,123,338 |
| LA50 | 196 |
| LGA50 | 211 |
| LA75 | 473 |
| LGA75 | 543 |

*All statistics are based on contigs of size >= 500 bp, unless otherwise noted (e.g., "# contigs (>= 0 bp)" and "Total length (>= 0 bp)" include all contigs).

**Supplementary Table 7. QUAST[29] analysis of *de novo* assemblies of NA12878 from different linked read methods[12-14] and nanopore method[26]. Assembly results from other linked read methods and nanopore long read method were cited from Supplemental Table S5 in Wang et al, 2019[14].**

| Assembly | TELL-Seq | stLFR-1[14] | stLFR-2[14] | 10x[14] | Nanopore[14] |
|---|---|---|---|---|---|
| # contigs (>= 0 bp) | 34,085 | 18,102 | 18,905 | 17,016 | 2,337 |
| # contigs (>= 25000 bp) | 1,181 | 965 | 948 | 697 | 1,819 |
| # contigs (>= 50000 bp) | 542 | 500 | 500 | 351 | 1,332 |
| Total length (>= 0 bp) | 2,979,445,840 | 2,899,386,838 | 2,893,558,199 | 2,867,683,417 | 2,866,880,913 |
| Total length (>= 25000 bp) | 2,794,100,219 | 2,816,263,478 | 2,812,836,296 | 2,804,290,035 | 2,858,326,432 |
| Total length (>= 50000 bp) | 2,772,210,571 | 2,800,220,687 | 2,797,538,549 | 2,792,294,955 | 2,840,893,649 |
| Largest contig | 109,183,970 | 104,958,599 | 86,747,726 | 109,375,584 | 50,410,306 |
| Total length | 2,979,445,840 | 2,848,642,889 | 2,836,789,635 | 2,824,111,036 | 2,866,386,869 |
| Reference length | 3,099,922,541 | 3,099,922,541 | 3,099,922,541 | 3,099,922,541 | 3,099,922,541 |
| GC (%) | 40.91 | 40.88 | 40.82 | 40.84 | 40.86 |
| Reference GC (%) | 40.87 | 40.87 | 40.87 | 40.87 | 40.87 |
| N50 | 31,462,027 | 28,877,588 | 26,308,501 | 45,263,628 | 7,667,013 |
| # misassemblies | 1,987 | 4,096 | 3,824 | 2,750 | 6,938 |
| Genome fraction (%) | 93.732 | 92.976 | 92.628 | 93.776 | 94.611 |
| # N's per 100 kbp | 6419.8 | 3039.36 | 2852.41 | 1085.71 | 0 |
| # mismatches per 100 kbp | 114.07 | 106.08 | 105.01 | 104.98 | 168.00 |
| # indels per 100 kbp | 25.61 | 25.64 | 25.56 | 28.32 | 165.25 |
| Largest alignment | 23,573,913 | 10,614,129 | 9,113,736 | 16,615,675 | 6,837,121 |
| NA50 | 4,302,918 | 1,863,160 | 1,983,491 | 2,863,578 | 906,345 |

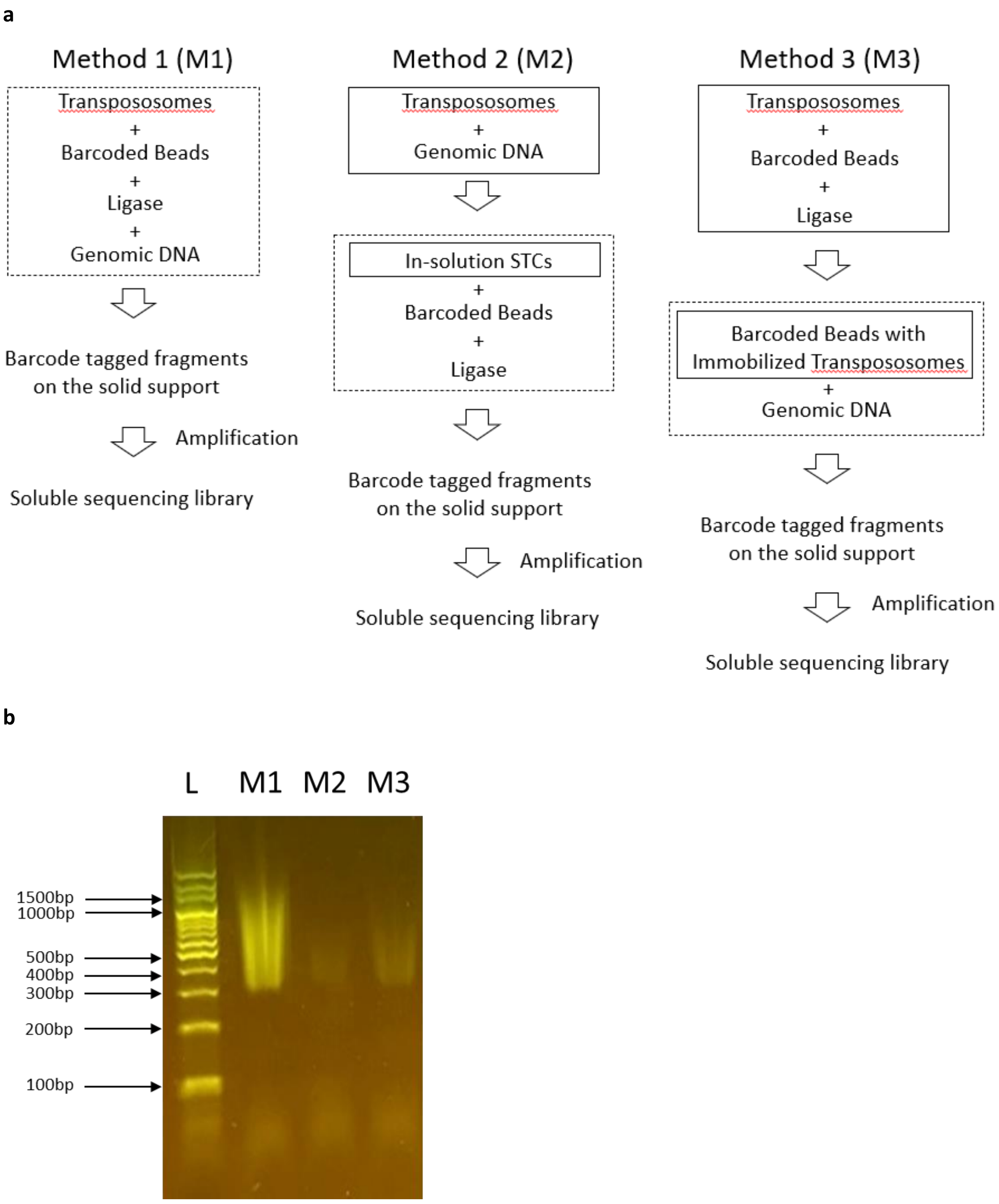


**Supplementary Figure 1**. Three different transposase based clonal barcoding methods without droplet facilitated compartmentation. (**a**) Overview of three different transposase based clonal barcoding methods. (**b**) An E-gel picture of amplified barcoded libraries using the three different transposase based clonal barcoding methods. L, 100bp DNA ladder; M1, Method 1; M2, Method 2; M3, Method 3. Expected library product size was greater than 300 bp.

Hg19: chr5:104432113-104503673 -> Hg38:chr5:105096412-105167972

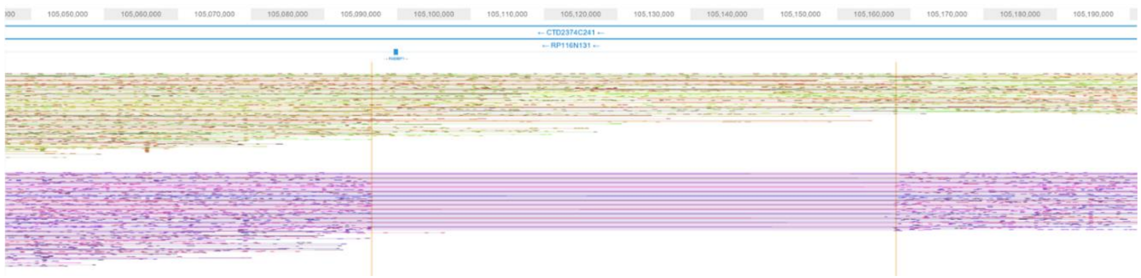

**Supplementary Figure 2**. Phased read graph from TELL-Seq data showed a heterozygous deletion on chromosome 5: 104432113-104503673. 10x data[12] called this region as a heterozygous deletion, while stLFR data[14] called this region as a homozygous deletion. Hg38 coordinates were used for visualization data.

Hg19: chr1:189704509-189783359 -> Hg38: chr1:189735379-189814229

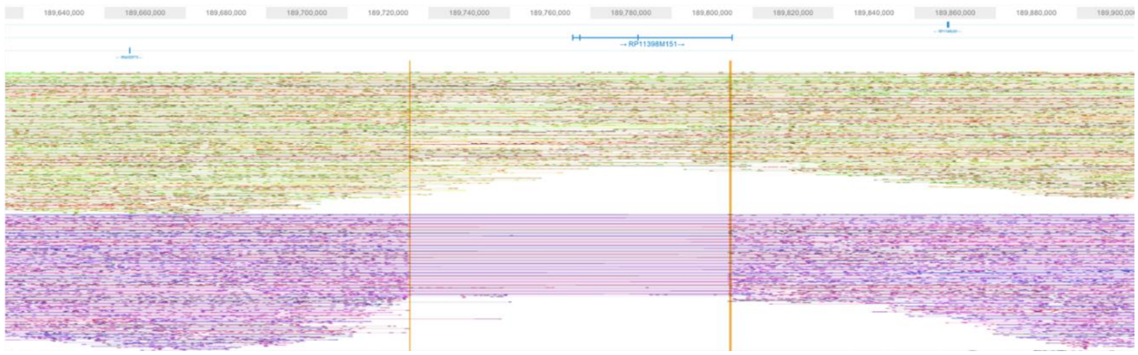

**Supplementary Figure 3**. Phased read graph from TELL-Seq data showed a heterozygous deletion on chromosome 1: 189704509-189783359. Hg38 coordinates were used for visualization data.

Hg19: chr3:65189000-65213999 -> Hg38: chr3:65203325-65228324

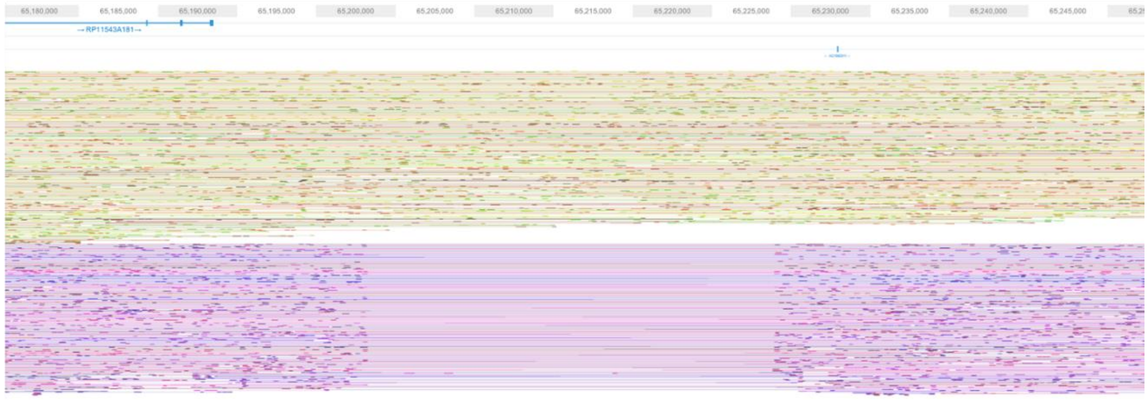

**Supplementary Figure 4**. Phased read graph from TELL-Seq data showed a heterozygous deletion on chromosome 3: 65189000-65213999. Hg38 coordinates were used for visualization data.

Hg19: chr4:116167000-116176999 -> Hg38: chr4:115245844-115255843

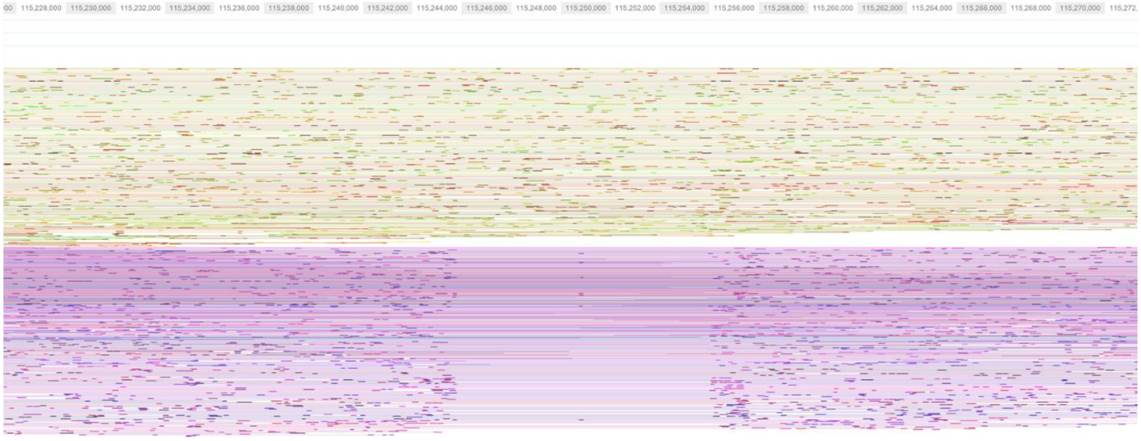

**Supplementary Figure 5**. Phased read graph from TELL-Seq data showed a heterozygous deletion on chromosome 4: 116167000-116176999. Hg38 coordinates were used for visualization data.

Hg19: chr4:187094000-187097999 -> Hg38: chr4:186172846-186176845

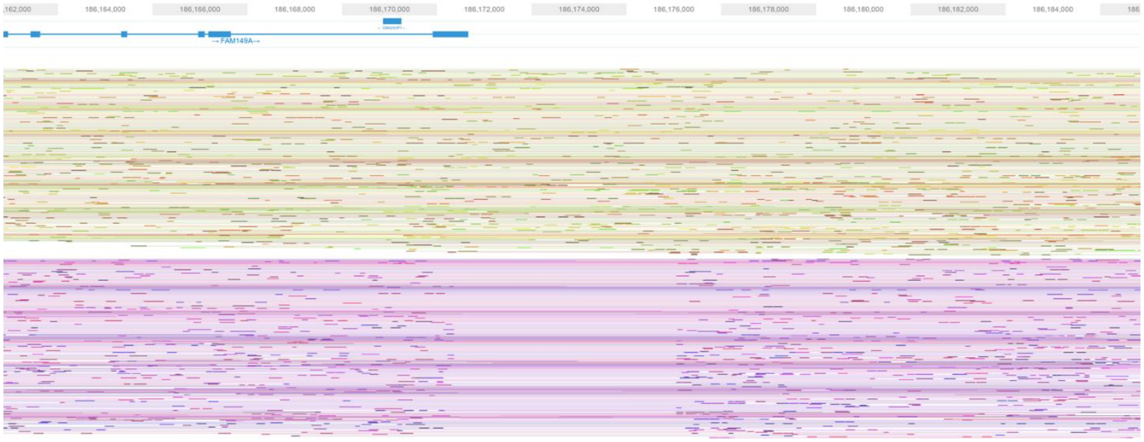

**Supplementary Figure 6**. Phased read graph from TELL-Seq data showed a heterozygous deletion on chromosome 4: 187094000-187097999. Hg38 coordinates were used for visualization data.

Hg19: chr6:78967194-79036419 -> Hg38: chr6:78257477-78326702

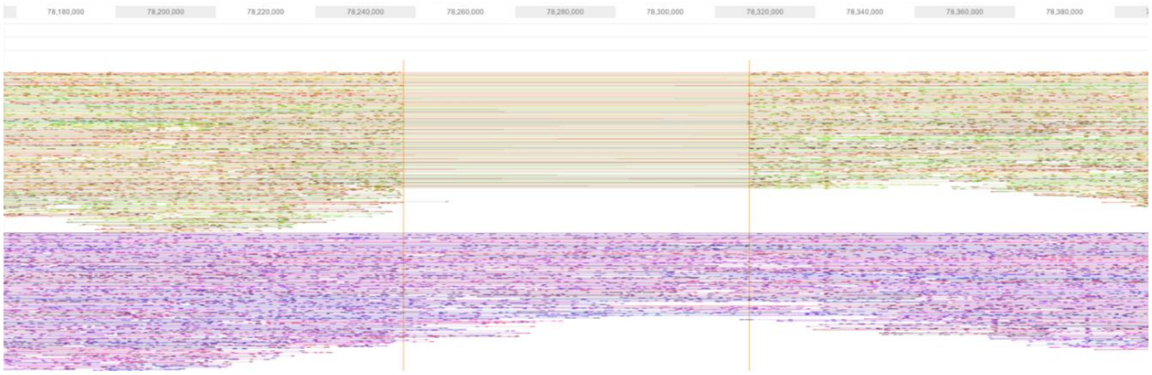


**Supplementary Figure 7**. Phased read graph from TELL-Seq data showed a heterozygous deletion on chromosome 6: 78967194-79036419. Hg38 coordinates were used for visualization data.

Hg19: chr7:110182000-110187999 -> Hg38: chr7:110541943-110547942

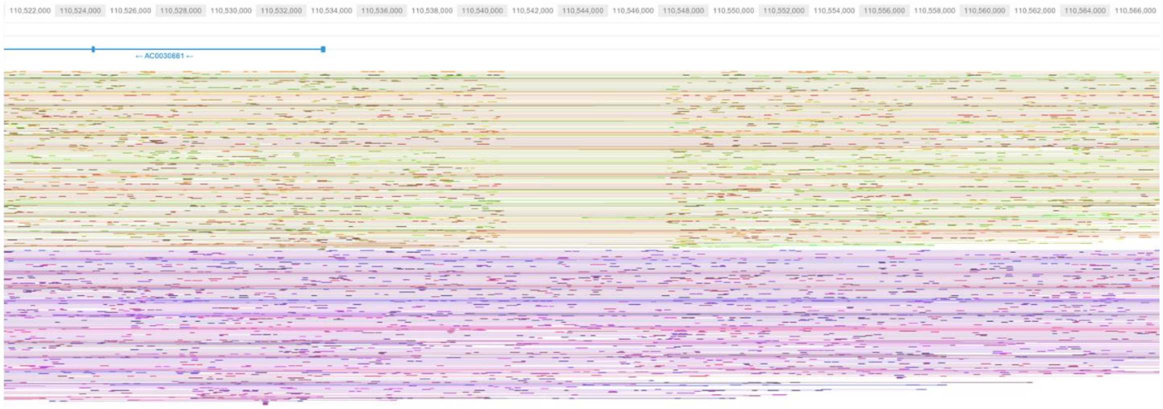


**Supplementary Figure 8**. Phased read graph from TELL-Seq data showed a heterozygous deletion on chromosome 7: 110182000-110187999. Hg38 coordinates were used for visualization data.

Hg19: chr8:39232074-39387229 -> Hg38: chr8:39374555-39529710

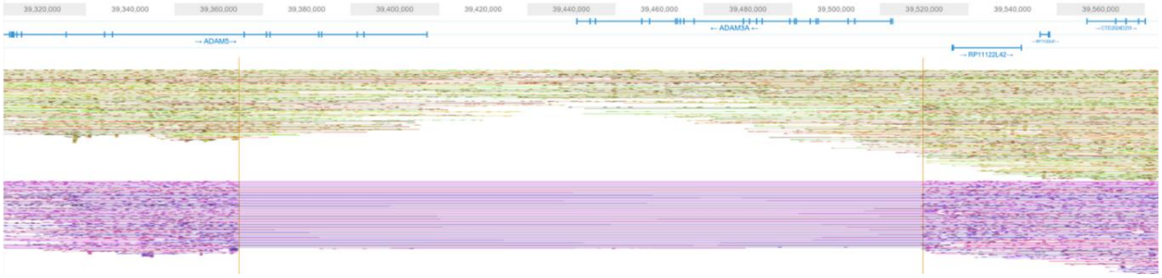


**Supplementary Figure 9**. Phased read graph from TELL-Seq data showed a heterozygous deletion on chromosome 8: 39232074-39387229. Hg38 coordinates were used for visualization data.

Hg19: chr16:62545000-62549999 -> Hg38: chr16:62511096-62516095

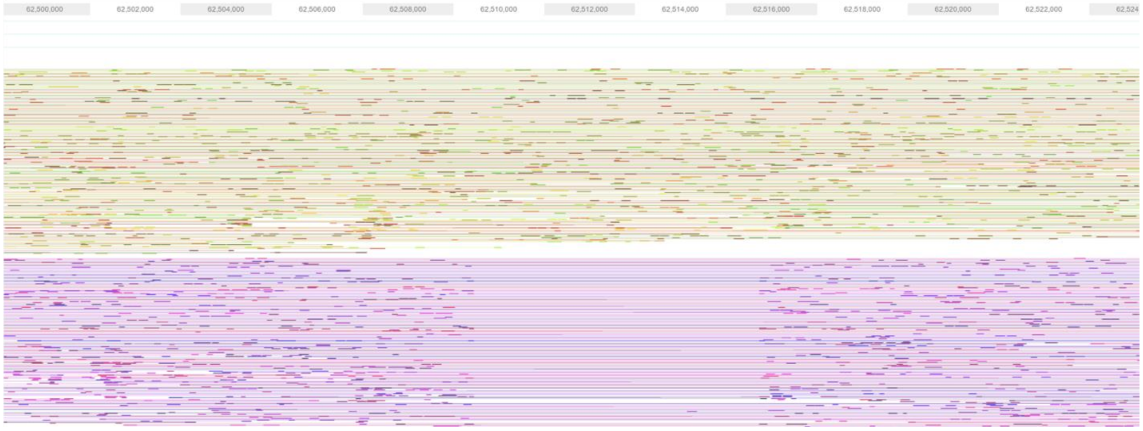


**Supplementary Figure 10**. Phased read graph from TELL-Seq data showed a heterozygous deletion on chromosome 16: 62545000-62549999. Hg38 coordinates were used for visualization data.
